# A daily-updated database and tools for comprehensive SARS-CoV-2 mutation-annotated trees

**DOI:** 10.1101/2021.04.03.438321

**Authors:** Jakob McBroome, Bryan Thornlow, Angie S. Hinrichs, Nicola De Maio, Nick Goldman, David Haussler, Russell Corbett-Detig, Yatish Turakhia

**Author notes:** Equal contribution (Jakob McBroome, Bryan Thornlow).

## Abstract

The vast scale of SARS-CoV-2 sequencing data has made it increasingly challenging to comprehensively analyze all available data using existing tools and file formats. To address this, we present a database of SARS-CoV-2 phylogenetic trees inferred with unrestricted public sequences, which we update daily to incorporate new sequences. Our database uses the recently-proposed mutation-annotated tree (MAT) format to efficiently encode the tree with branches labeled with parsimony-inferred mutations as well as Nextstrain clade and Pango lineage labels at clade roots. As of June 9, 2021, our SARS-CoV-2 MAT consists of 834,521 sequences and provides a comprehensive view of the virus’ evolutionary history using public data. We also present matUtils – a command-line utility for rapidly querying, interpreting and manipulating the MATs. Our daily-updated SARS-CoV-2 MAT database and matUtils software are available at http://hgdownload.soe.ucsc.edu/goldenPath/wuhCor1/UShER_SARS-CoV-2/ and https://github.com/yatisht/usher, respectively.

## Introduction

The COVID-19 pandemic has witnessed unprecedented levels of genome sequencing for a single pathogen (Hodcroft et al. 2021). Since the onset of the pandemic in late 2019, over a million SARS-CoV-2 genomes have been sequenced worldwide, and tens of thousands of new genomes are being shared on various data repositories every day (Maxmen 2021). This data has enabled scientists to closely track the evolution of the virus and study its transmission dynamics at global and local scales (Deng et al. 2020; Chaillon and Smith 2021; da Silva Filipe et al. 2021). However, the scale of this data is posing serious computational challenges for comprehensive phylogenetic analyses (Hodcroft et al. 2021). Platforms like Nextstrain (Hadfield et al. 2018) have been invaluable in studying viral transmission networks and genomic surveillance efforts, but they only provide subsampled SARS-CoV-2 trees consisting of a tiny fraction of available data, omitting phylogenetic relationships with most available sequences. A single, comprehensive SARS-CoV-2 reference tree of all available data could not only facilitate detailed and unambiguous phylogenetic analyses at global, country and local levels, but may also help promote consistency of results across different research groups (Turakhia et al. 2020).

Besides the computational challenges, the massive volume of SARS-CoV-2 data is also posing numerous data sharing challenges with existing file formats, such as Fasta or Variant Call Format (VCF), which are bulky and necessitate network speeds and computational capabilities that are beyond the reach of many research and scientific groups involved in studying SARS-CoV-2 evolution and transmission dynamics worldwide.

## New Approaches

In this work, we simultaneously address the issue of maintaining a comprehensive SARS-CoV-2 reference tree and its associated data processing, data sharing and computational analysis challenges. Specifically, we are maintaining and openly sharing a daily-updated database of mutation-annotated trees (MATs) containing global SARS-CoV-2 sequences from public databases, including annotations for Nextstrain clades (Hadfield et al. 2018) and Pango lineages (Rambaut et al. 2020) (Supplementary Figure 1). The MAT is an extremely efficient data format proposed recently (Turakhia et al. 2021) which can facilitate the sharing of extremely large genome sequence datasets – an uncompressed MAT of 834,521 SARS-CoV-2 public sequences requires only 65 MB to store, and encodes more information than is contained in a 43 GB VCF and a 38 MB Newick file combined.

To accompany this database, we present matUtils – a toolkit for querying, interpreting and manipulating the MATs. Using matUtils, common operations in genomic surveillance and contact tracing efforts, including annotating a MAT with new clades, extracting subtrees of the most closely-related samples, or converting the MAT to standard Newick or VCF format can be performed in a matter of seconds to minutes even on a laptop. We also provide a web interface for matUtils through the UCSC SARS-CoV-2 Genome Browser (Fernandes et al. 2020). Together, our SARS-CoV-2 database and matUtils toolkit can simultaneously democratize and accelerate pandemic-related research.

## Results and Discussion

### A daily-updated mutation-annotated tree database of global SARS-CoV-2 sequences

To aid the scientific community studying the mutational and transmission dynamics of the SARS-CoV-2 virus and its different variants, we are maintaining a daily-updated database of SARS-CoV-2 mutation-annotated trees (MATs) composed of public data. Starting with the final Newick tree release dated November 13, 2020, of Rob Lanfear’s sarscov2phylo (https://github.com/roblanf/sarscov2phylo) that is re-rooted to Wuhan/Hu-1 (GenBank MN908947.3, RefSeq NC_045512.2), we have set up an automated pipeline to aggregate public sequences available through GenBank (Clark et al. 2007), COG-UK (Nicholls et al. 2020), and the China National Center for Bioinformation on a daily basis and incorporate them into our MAT using UShER (see Supplementary Methods). GISAID data (Shu and McCauley 2017) is not included in our MATs because its usage terms do not allow redistribution. We also use the *matUtils annotate* command (see Supplementary Methods) to add Nextstrain clade and Pango lineage annotations to individual branches of our MAT. As of June 9, 2021, our MAT consists of 834,521 sequences, includes 14 Nextstrain clade and 895 Pango lineage annotations for all samples, and is only 65 MB, or 14 MB in its gzip-compressed form (Supplementary Figure 1, Supplementary Table S1). To our knowledge, this is the most comprehensive representation of the SARS-CoV-2 evolutionary history using publicly available sequences. It can be freely used to study evolutionary and transmission dynamics of the virus at global, country and local levels.

### matUtils provides a wide range of functions to analyze and manipulate mutation-annotated trees

We have created a high-performance command line utility called matUtils for performing a wide range of operations on MATs for rapid interpretation and analysis in genomic surveillance and contact tracing efforts. matUtils is distributed with the UShER package (Turakhia et al. 2021) and uses the same mutation-annotated tree (MAT) format as UShER. matUtils is organized into five different subcommands: annotate, summary, extract, uncertainty and introduce (Figure 1), described briefly below. We provide detailed instructions for the usage of each module on our wiki (https://usher-wiki.readthedocs.io/en/latest/matUtils.html).

**Figure 1:**
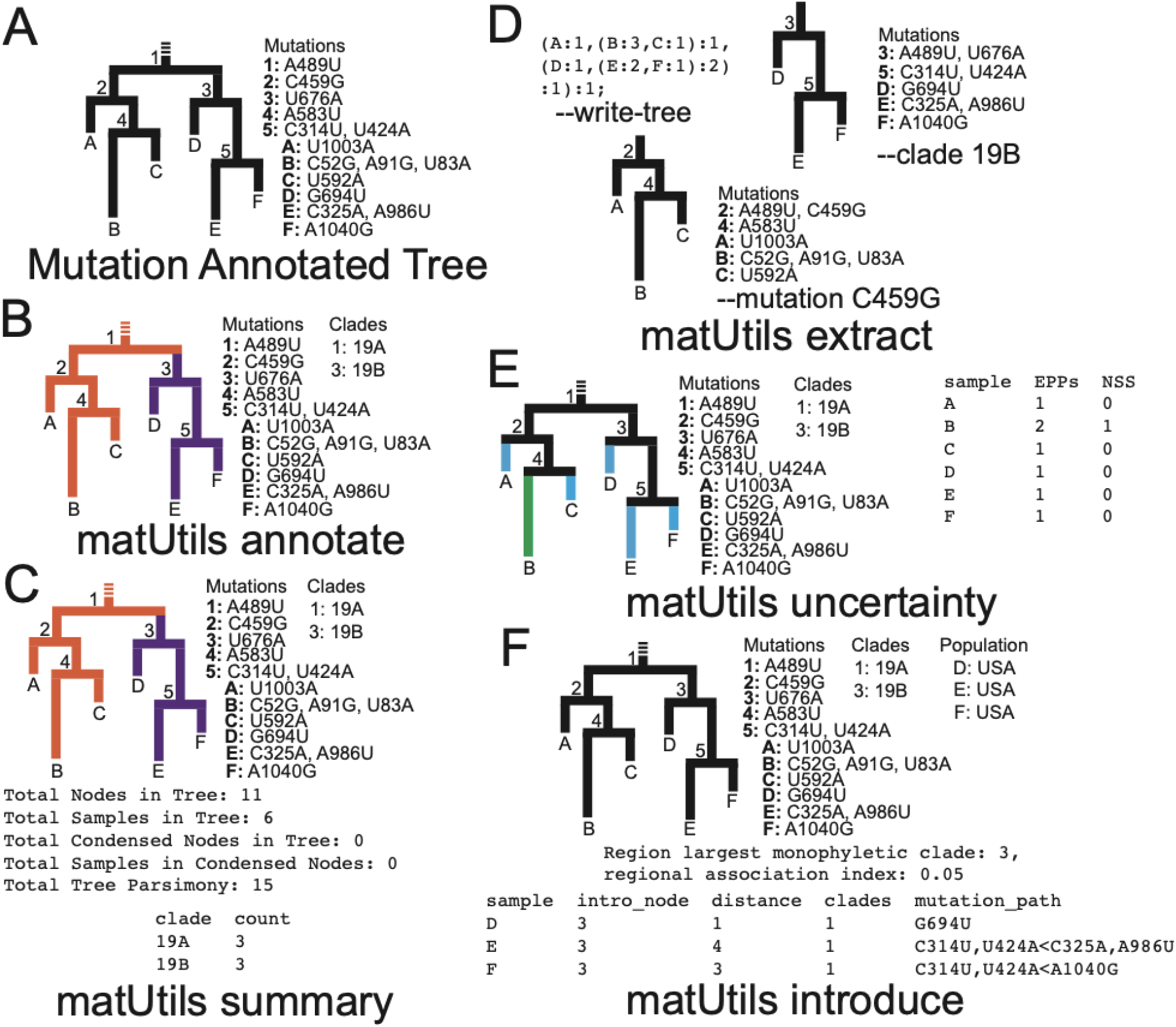
matUtils functions enable fast, user-friendly analysis of mutation-annotated trees (MATs). **(A)**: An example MAT with tree topology corresponding to the MAT on the left and the mutation annotations on each node shown on the right. **(B):** *matUtils annotate* allows the user to annotate internal nodes with clade names. In this example, nodes 1 and 3 are annotated with clade names 19A and 19B, respectively. **(C):** *matUtils summary* outputs sample-, clade-, and tree-level statistics for the input MAT. **(D):** *matUtils extract* allows users to convert a MAT (of panel C in this example) to Newick format (left), subset the MAT for a specified clade (right) or mutation (bottom), among other functions. **(E)**: *matUtils uncertainty* outputs parsimony scores, equally parsimonious placements (EPPs) and neighborhood size scores (NSS) for each sample. **(F):** *matUtils introduce* takes as input a list of samples of interest, and outputs their predicted introduction nodes and paths, along with confidence scores for both the introduction and parent node (not shown). Using the *-a* flag, the user can also determine the largest monophyletic clade and regional association index associated with the input population. Where relevant for all functions shown, text outputs are displayed in fixed-width fonts for differentiation from attributes internal to the input protobuf.

#### Annotate

This function provides the ability to annotate the clades in the tree. One of the central uses of phylogenetics during the pandemic is to trace the emergence and spread of new viral lineages. Nextstrain (Hadfield et al. 2018), Pango (Rambaut et al. 2020) and GISAID (Shu and McCauley 2017) provide different nomenclatures for SARS-CoV-2 variants that have been used widely in genomic surveillance. Our MAT format provides the ability to annotate internal branches of the tree with an array of clade names, one for each clade nomenclature. Clades can be annotated on a MAT using *matUtils annotate* in two ways: (i) directly providing the mappings of each clade name to its corresponding node or (ii) providing a set of representative sample names for each clade from which the clade roots can be automatically inferred (see Supplementary Methods). Both ways of annotating ensure that the clades remain monophyletic, but we use the second approach to label Nextstrain clades and Pango lineages in our SARS-CoV-2 MAT database since it can be automated using available data (see Supplementary Methods). *matUtils annotate* has high congruence with Nextstrain clades and Pango lineage annotations (Supplementary Table S1).

Once clades are annotated on a MAT, the UShER placement tool (Turakhia et al. 2021) can assign each newly placed sequence to its corresponding Pango lineage, and this being used as a feature in Pangolin 3.0 (https://github.com/cov-lineages/pangolin/releases/tag/v3.0) to perform clade assignments in a fully phylogenetic framework.

#### Summary

This function provides a brief summary of the available data in the input MAT file and is meant to serve as a typical first step in any MAT-based analysis. It provides a count of the total number of samples in the MAT, the number of annotated clades, the size of each clade, the total parsimony score (i.e. the sum of mutation events on all branches of the MAT), the number of distinct mutations, clade assignments for each sample, and other similar statistics.

#### Extract

Many SARS-CoV-2 phylodynamic studies involve restricting the analysis to a smaller tree of interest, such as a tree of sequences belonging to a particular geographic region or clade. It can be computationally challenging to identify samples most closely related to a given sample or cluster from over a million other sequences, or infer individual subtrees, but it is straightforward to retrieve subtrees from a comprehensive phylogeny. *matUtils extract* provides an efficient and robust suite of options for subtree selection from a MAT that could transform viral genomic surveillance efforts. A user can use *matUtils extract* to subsample a MAT to find samples that contain a mutation of interest, are members of a specific clade, have a name matching a specific regular expression pattern (such as the expression “(IND*|India*)” to select samples from India), among other criteria (see Supplementary Methods). *matUtils extract* also includes options for pruning low-quality sequences from a MAT, such as those with an unusually high parsimony score. Notably, *matUtils extract* can produce an output Auspice v2 JSON that is compatible with the Auspice tree visualization tool (Hadfield et al. 2018) (Figure 2, Supplementary Methods). *matUtils extract* can also convert a MAT into other file formats, such as a Newick for its corresponding phylogenetic tree and a VCF for its corresponding genome variation data.

**Figure 2:**
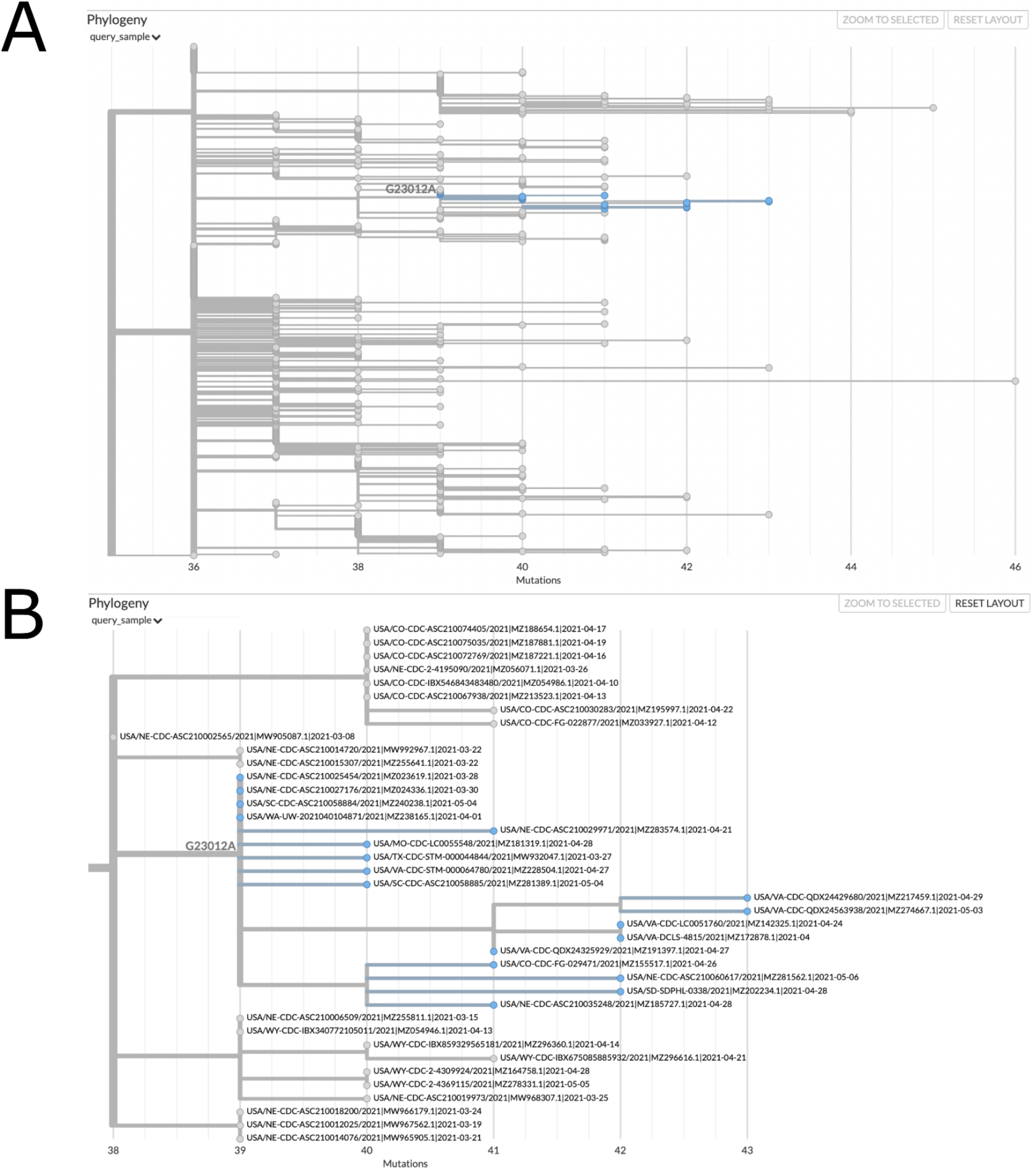
matUtils can generate informative visuals with Auspice. The above trees represent a clade of related B.1.1.7 samples from the USA which secondarily acquired the potentially important spike protein mutation E484K, which is caused by the nucleotide mutation G23012A. These trees were obtained by running the command “matUtils extract -i public-2021-06-09.all.masked.nextclade.pangolin.pb.gz -c B.1.1.7 -m G23012A -H “(USA.*)” -N 500 -j clade_trees -d clade_out”, which selects all samples from clade B.1.1.7 which acquired this mutation and are from the USA, then identifies the minimum set of five hundred sample subtrees which contain all of these samples, creating an Auspice v2 format JSON for each subtree (Hadfield et al 2018). This results in thirty-five distinct subtree JSON files of five hundred samples each in the output directory. Panel A represents the entirety of subtree six as viewed with Auspice (Hadfield et al 2018), including blue highlights and a branch label where our mutation of interest occurred. Panel B is zoomed in on this subtree and its sister clade; at this scale we can read individual sample names and observe that this specific strain has been actively spreading in the United States during April 2021.

#### Uncertainty

A fundamental concern in SARS-CoV-2 phylogenetics is topological uncertainty (Hodcroft et al. 2021). This is especially true for public health, where sample level uncertainty statistics convey the reliability of genomic contract tracing. matUtils provides such a statistic through its uncertainty function, which computes the number of equally parsimonious placements (Turakhia et al. 2020) that exist for each specified sample in the input MAT. Importantly, matUtils also allows the user to calculate equally parsimonious positions for already placed samples. This is accomplished by pruning the sample from the tree and placing the sample back to the tree using the placement module of UShER (Turakhia et al. 2021) (see Supplementary Methods). *matUtils uncertainty* additionally records the number of mutations separating the two most distant equally parsimonious placements, reflecting the distribution of placements across the tree (see Supplementary Methods). The output file is compatible as “drag-and-drop” metadata with the Auspice platform which allows for a rapid visualization of sample placement uncertainty (Supplementary Figure 2)

#### Introduce

Public health officials are often concerned about the number of new introductions of the virus genome in a given country or local area. To aid this analysis, *matUtils introduce* can calculate the association index (Wang et al. 2001) or the maximum monophyletic clade size statistic (Salemi et al. 2005; Parker et al. 2008) for arbitrary sets of samples, along with simple heuristics for approximating points of introduction into a region (see Supplementary Methods).

### matUtils enables rapid analysis of a comprehensive SARS-CoV-2 global tree and its web interface

The sheer scale of genomic data collected during the ongoing SARS-CoV-2 pandemic has necessitated the development of new tools for effective phylogenetic analysis. The matUtils toolkit is designed to scale efficiently to SARS-CoV-2 phylogenies containing millions of samples. Using matUtils, common pandemic-relevant operations described in the earlier section can be performed in the order of seconds to minutes with the current scale of SARS-CoV-2 data (Supplementary Tables S2-S9). For example, it takes only 5 seconds to summarize the information contained in our 06/09/2021 SARS-CoV-2 MAT of 834,521 samples and only 15 seconds to extract the mutation paths from the root to every sample in the MAT (Supplementary Table S2). Since matUtils is primarily designed to work with the newly-proposed and information-rich MAT format, it does not have direct counterparts in other bioinformatic software packages currently, but its efficiency is similar or better than state-of-the-art tools that offer comparable functionality (Supplementary Tables S2-S9). For example, matUtils is able to resolve polytomies in a 834,521 sample tree in 9 seconds, a task which takes over 37 minutes using ape (Paradis and Schliep 2019) (Supplementary Table S3). matUtils is also very memory-efficient, requiring less than 1.4 GB of main memory for most tasks, making it possible to run even on laptop devices.

Certain functions of matUtils (such as extracting subtrees of provided sample names or identifiers) have also been ported to UCSC SARS-CoV-2 Genome Browser (Fernandes et al.2020) and are available from https://genome.ucsc.edu/cgi-bin/hgPhyloPlace. This provides a user-friendly web interface to public health officials and researchers working on combating the pandemic.

Our database and utility fill a critical need for open, public, rapid analysis of the global SARS-CoV-2 phylogeny by health departments and research groups across the world, with highly-efficient file formats that do not require high speed internet connectivity or large storage devices, and tools capable of rapidly performing large-scale analyses on laptops.

## Acknowledgments

We thank Rob Lanfear for reviewing this manuscript and his valuable feedback. We thank Cheng Ye for his help in parallelizing VCF extraction. We also thank all the laboratories that submit data to public databases.

## Supplementary Methods

### Maintaining a daily-updated mutation-annotated tree database of global SARS-CoV-2 sequences

We are maintaining a daily-updated mutation-annotated tree (MAT) database of global SARS-CoV-2 sequences at http://hgdownload.soe.ucsc.edu/goldenPath/wuhCor1/UShER_SARS-CoV-2/. Our database is organized into sub-directories sorted by year, month and date. To update the MATs daily, we have set up a CRON job on a server at UCSC which downloads SARS-CoV-2 sequences daily from GenBank (Clark et al. 2007) and COG-UK (Nicholls et al. 2020) (see https://github.com/ucscGenomeBrowser/kent/blob/master/src/hg/utils/otto/sarscov2phylo/updatePublic.sh which calls other scripts in the same directory). We also include 253 sequences downloaded from the China National Center for Bioinformation (https://bigd.big.ac.cn/ncov/release_genome) in October 2020 that are not associated with GenBank IDs.

New sequences are added to the previous day’s MAT using the UShER placement tool (Turakhia et al. 2021) with options to place the samples in the order of the fewest ambiguous bases and exclude sequences with 5 or more equally parsimonious placements. Previously excluded sequences are reconsidered for placement during each build. We also use *matUtils extract* to prune samples with 30 or more private mutations and those internal branches longer than 30 mutations, as these are highly indicative of error-containing sequences (Mai and Mirarab 2018). The trees are rooted to Wuhan/Hu-1 (GenBank MN908947.3, RefSeq NC_045512.2), and nodes with no associated mutations are collapsed (Turakhia et al. 2021). Our first MAT was created by starting with the last Newick tree release (dated November 13, 2020) of Rob Lanfear’s sarscov2phylo (Lanfear and Mansfield 2020) containing 82,358 public sequences, adding the later additional public sequences using UShER. Each MAT is then annotated with Nextstrain clade and Pango lineage annotations using *matUtils annotate -c* with a file containing representative sequences for each clade/lineage. For Nextstrain clades, Nextclade assignments (https://github.com/nextstrain/nextclade) for all sequences are used. For Pango lineages, designated lineage representative sequences from https://github.com/cov-lineages/pango-designation/ are mapped to the corresponding public sequence IDs where possible.

In addition to MATs, we provide in each sub-directory: (i) a Variant Call Format (VCF) file containing the genotypes of public sequences, generated from the corresponding MAT with matUtils extract such that missing or ambiguous bases have been imputed by UShER using maximum parsimony (Turakhia et al. 2021), (ii) a Newick file also generated from the corresponding MAT using *matUtils extract* (iii) a tab-separated file containing information about each public sequence e.g. collection date, location, Nextstrain clade and Pango lineage, (iv) a tab-separated file with Nextstrain clades assigned to sequences by Nextclade (https://github.com/nextstrain/nextclade) and (v) a tab-separated file with Pango lineages assigned to sequences by pangolin (https://github.com/cov-lineages/pangolin).

Our script to update the MAT daily is available at https://github.com/ucscGenomeBrowser/kent/blob/master/src/hg/utils/otto/sarscov2phylo/updateCombinedTree.sh.

### matUtils: Design Overview

matUtils is implemented using the C++ programming language and is developed and maintained within the phylogenetic placement package of UShER (Turakhia et al. 2021), since matUtils shares the core mutation-annotated tree (MAT) data structure with UShER, which helps us ensure cross-compatibility of both tools. matUtils complements UShER through its ability to analyze and manipulate the MAT output but can be used as a standalone phylogenetics tool independent of UShER. matUtils and UShER can be installed together via (i) a Docker container (https://hub.docker.com/repository/docker/yatisht/usher), (ii) the Conda package manager using the bioconda (Grüning et al. 2018) channel (http://bioconda.github.io/recipes/usher/README.html) or (iii) the installation scripts that we provide on our GitHub repository (https://github.com/yatisht/usher) for some recent Linux and MacOS releases. Detailed installation and usage instructions are available on our wiki: https://usher-wiki.readthedocs.io/en/latest/matUtils.html. Several matUtils functions have multi-threaded parallel implementations through Intel’s Thread Building Blocks library (https://github.com/oneapi-src/oneTBB).

### matUtils: Implementation details

#### Annotate

*matUtils annotate* is designed to annotate clades on the internal branches of the MAT. Our MAT format (specified in https://github.com/yatisht/usher/blob/master/parsimony.proto) provides an ability to annotate internal branches with an array of clade names, one for each clade nomenclature. Each run of *matUtils annotate* extends the clade name array size in a MAT by one to accommodate a new nomenclature. Only the node corresponding to a clade root is labeled with its clade name, as descendants of that node can be automatically inferred to belong to that clade. Clades can be nested, so that each sequence can be assigned to a clade corresponding to the lowest-level clade root to which it is a descendant. *matUtils annotate* provides two different ways to annotate clades in a MAT. Both ways, by design, ensure that all clades remain monophyletic. In the first, a user can directly provide the internal node identifiers corresponding to the root of each clade. In the second, a user can provide a list of representative sequences for each clade, such as training data for Pango lineages (https://github.com/cov-lineages/pango-designation), from which the clade root can be inferred in the tree. Not all sequences in the tree need to be designated by a clade. Since the training data is imperfect, and the representative sequences for lineages are sometimes non-monophyletic in our tree, we have found the simple approach of using the most recent common ancestor (MRCA) does not yield accurate results. The *matUtils annotate* inference method works instead by first building a “consensus” sequence (where, by default, the consensus sequence requires an allele to be present in at least 80% of representative sequences, with lower frequency alleles marked as ambiguous) for each clade and finding its phylogenetic placement using UShER’s placement module to obtain the clade root. When multiple equally parsimonious placements are available for the clade root, the algorithm uses a heuristic formula to compute the “best fit” for the training data, which rewards the placement containing a higher proportion of samples designated by that clade in the training data and penalizes descendants designated by some other clade in the training data. When the same root is found for multiple clades, the clade with fewest equally parsimonious placements, followed by the number of representative sequences in the training data, is prioritized.

#### Extract

The *extract* subcommand acts as a simple prebuilt pipeline with three distinct stages. The first of these, sample selection, collects the set of samples which fulfill each of the conditions indicated by input parameters, then gets the intersection of these sets to identify samples which fulfill all conditions specified on the command. Multiple conditions can be simultaneously specified in a single command for selecting samples, such as clade membership, maximum parsimony score, presence of a particular mutation, and whether the sample name matches a specific regular expression pattern, among others. The second stage edits the input tree object to generate the indicated subtree, either by pruning excluded samples or by generating a subtree in a parallelized fashion, depending on the size of the chosen sample input. The third stage generates each of the requested output files representing the final tree. These files include Newick for pure tree information, parsimony-resolved VCF for variation information, and Auspice v2 format JSON for both (Hadfield et al. 2018). VCF production is parallelized for efficiency with large sample selections. A sample metadata table in CSV or TSV format can be incorporated into the JSON output. The full list of options can be found at our wiki: https://usher-wiki.readthedocs.io/en/latest/matUtils.html.

#### Uncertainty

*matUtils uncertainty* can calculate two different metrics for characterizing the phylogenetic certainty of a sample placement. The first metric is “equally parsimonious placements” (EPPs), which is the number of places on the tree a sample could be placed without affecting the parsimony score. An EPP score of 1 indicates a high placement certainty of a sample in its local neighborhood in that there is a single most parsimonious placement location for that sample on the entire tree, and a higher EPP score suggests the sample placement is less certain. This metric is calculated by computing the number of most parsimonious placements after remapping the input sample(s) against the same tree (disallowing it from mapping to itself) with UShER’s optimized placement module. About 85% of samples in our SARS-COV-2 MAT database have an EPP score of 1. The second metric is “neighborhood size score” (NSS), which is the longest distance (in number of edges) between any two equally parsimonious placement locations for a given sample. This metric is complementary to EPPs – when multiple EPPs are possible for a sample, NSS indicates whether the placement uncertainty is restricted to a small neighborhood (small NSS value) or spans a large portion of the tree (large NSS value).

#### Introduce

*matUtils introduce* is aimed to help epidemiologists and public health officials estimate the number of new introductions of the virus in a given area or country. It includes a command which calculates maximum monophyletic clade size and association index statistics for phylogeographic trait association for user-provided input regions. Maximum monophyletic clade size (Parker et al. 2008) is the largest monophyletic clade of samples which are in the region – it is larger for regions which have relatively fewer introductions per sample and correlates with overall sample size. Association index (Wang et al. 2001) is a more complex metric which performs a weighted summation across the tree accounting for the number of child nodes and the frequency of the most common trait, such as membership in a particular geographical region of interest. Association index is smaller for stronger phylogeographic association and increases with the relative number of introductions into a region. For association index, *matUtils introduce* also performs a series of permutations to establish an expected range of values for the random distribution of samples across the tree. We are also working with epidemiologists currently to expand *matUtils introduce* with new heuristics (currently in an experimental stage but described in more detail on our wiki at: https://usher-wiki.readthedocs.io/en/latest/matUtils.html) for estimating the number of new introductions of the virus in a given area, details of which will appear in a later publication.

### Performance benchmarking of matUtils and other phylogenetics software packages

All performance benchmarking experiments were carried out on a Google Cloud Platform (GCP) instance n2d-standard-16 with 16 vCPUs (Intel Xeon CPU E7-8870 v.4, 2.10 GHz) with 64 GB of memory using our public SARS-CoV-2 MAT dated June 9, 2021. matUtils does not have direct counterparts for its ability to work with the mutation-annotated tree (MAT) format, but we compared the performance of matUtils with state-of-the-art tools that offer some comparable functionality on Newick or VCF formats. Specifically, we compared the most recent version of matUtils (version 0.3.1) to newick_utils version 1.6 (Junier and Zdobnov 2010), tree_doctor (from version 1.5 of the phast package; (Hubisz et al. 2011), ape version 5.5 (Paradis and Schliep 2019), and bcftools version 1.7 (Danecek et al. 2011). The exact commands used for each comparison can be found in Supplementary Tables S2-S9, and the input data used for each comparison can be found at https://github.com/bpt26/matutils_benchmarking/.

## Supplementary Figures and Tables

**Supplementary Figure 1:**
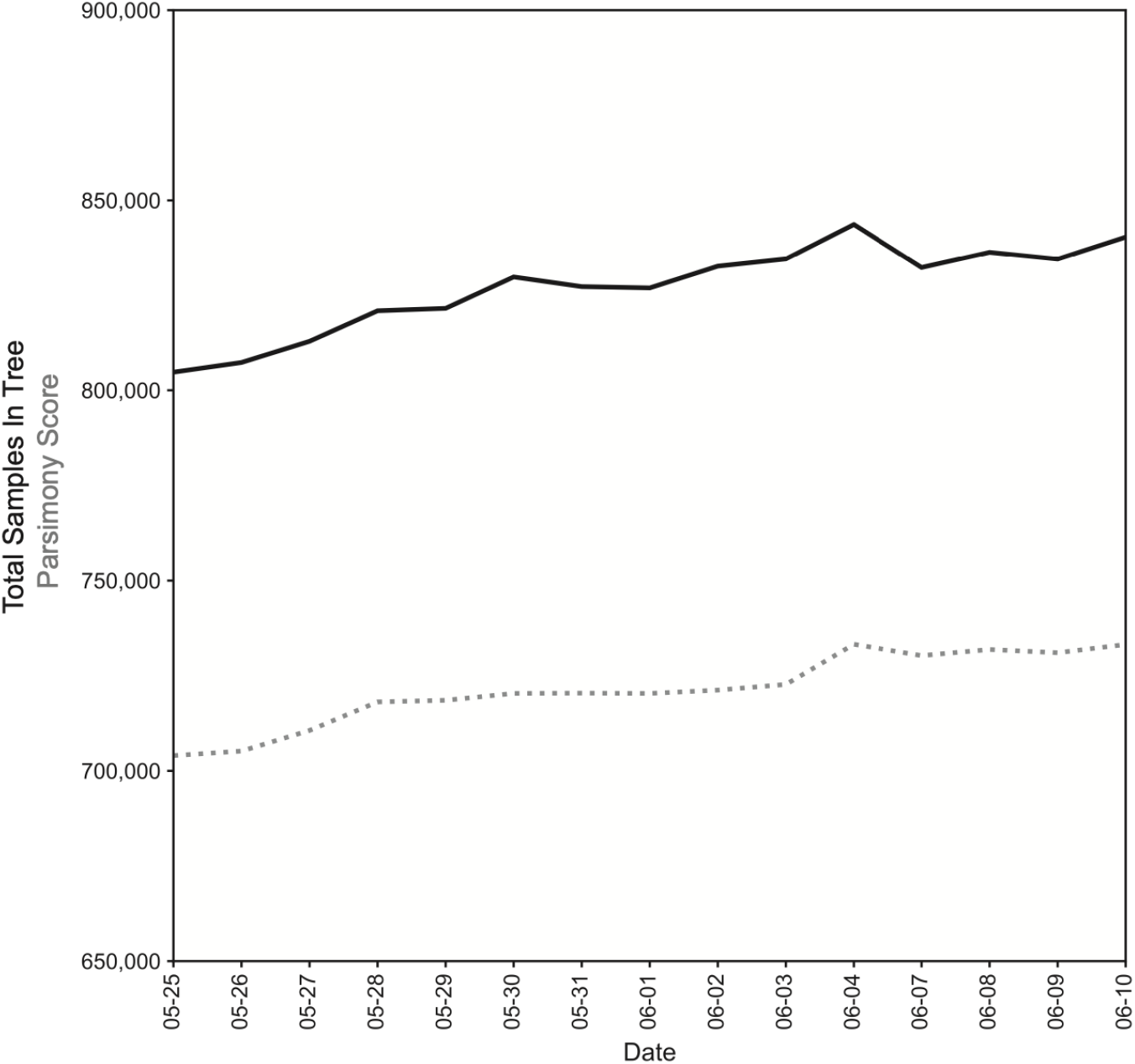
Our global phylogeny contains 840,343 samples as of June 10. Our database, containing all high-quality publically available SARS-CoV-2 whole-genome sequences and their clade assignments, is updated daily. As of June 10, our phylogeny contains 840,343 sequences with a total parsimony score of 733,211. Sequences that have 5 or more equally parsimonious placements on the tree are removed at each build (see Supplementary Methods), so the total samples sometimes drop during successive builds.

**Supplementary Figure 2:**
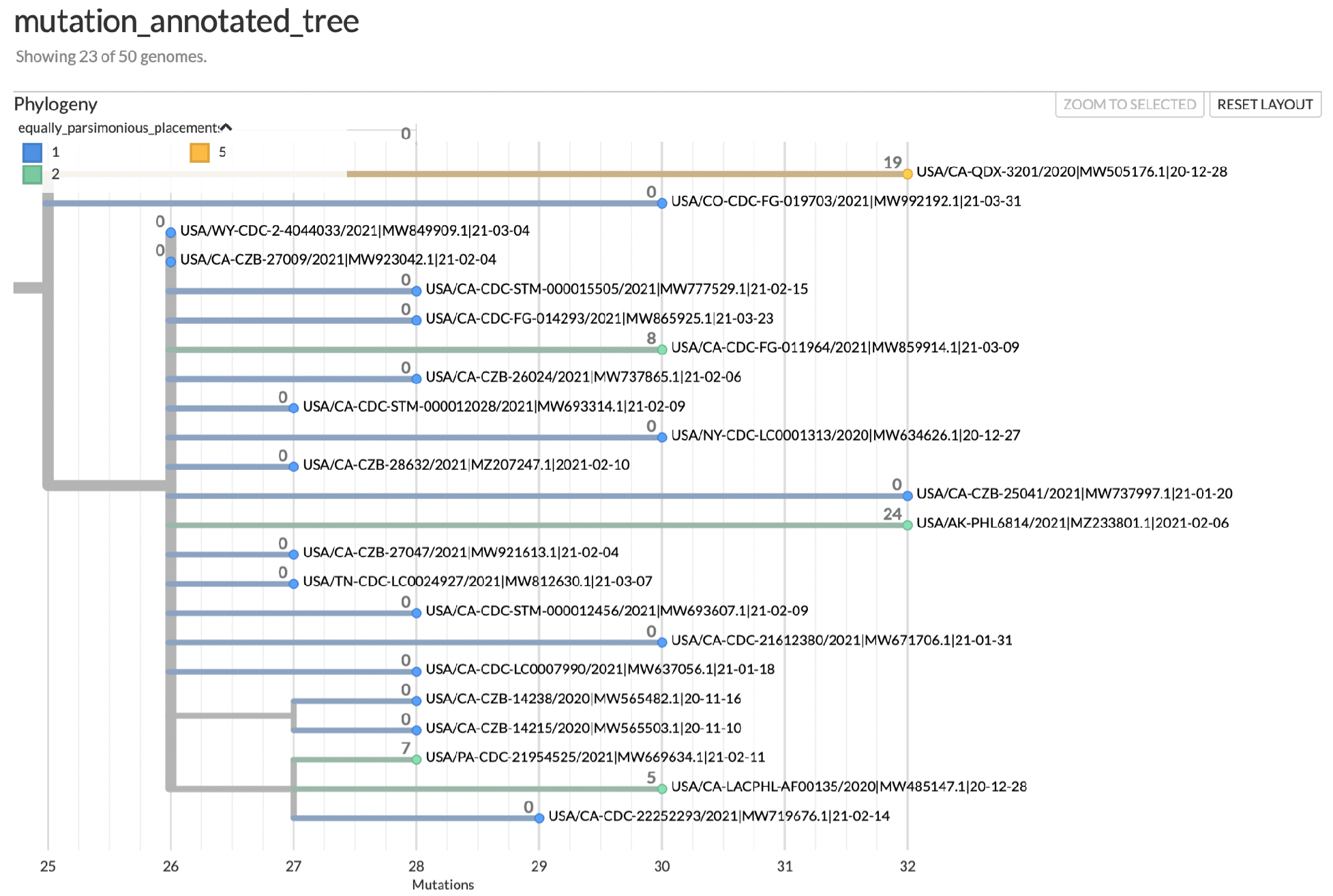
matUtils uncertainty statistics reveal low-quality sample placements. This Auspice view of an example subtree is annotated with both equally parsimonious placements (in color) and neighborhood size (branch label integers). 18 of our 23 samples in the subtree have a single placement and a neighborhood size of 0, indicating high placement certainty for those samples. Of the five samples with multiple equally parsimonious placements, one sample has 5 equally parsimonious placements with an NSS value of 19, indicating a high level of placement uncertainty for this sample spanning a relatively large neighborhood.

**Supplementary Table S1:**
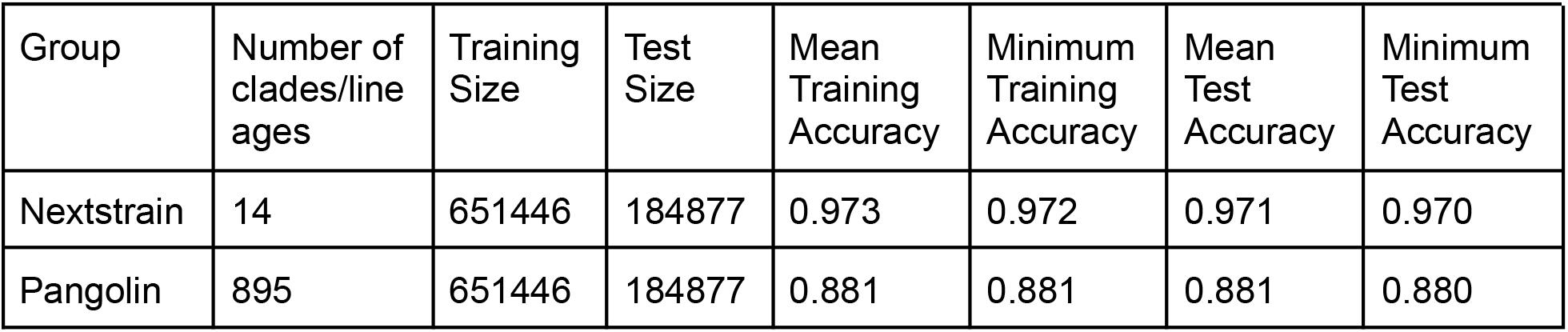
matUtils annotate can quickly and effectively assign clade lineage roots. This table was generated by taking training data associated with Nextstrain clades and Pango lineages from our public repository (lineageToPublicName.gz and cladeToPublicName.gz), splitting the data 80/20 into training and test sets, and assigning roots based on the 80% selected training data with matUtils annotate on the 06-09-2021 public MAT tree. Accuracy was scored as the percentage of the training or test set which matches Nextclade or Pangolin assignments. This process was repeated 9 times and mean and minimum accuracy values were collected.

## References

Chaillon A, Smith DM. 2021. Phylogenetic analyses of SARS-CoV-2 B.1.1.7 lineage suggest a single origin followed by multiple exportation events versus convergent evolution. Clinical Infectious Diseases [Internet]. Available from: https://doi.org/10.1093/cid/ciab265

Clark AG, Eisen MB, Smith DR, Bergman CM, Oliver B, Markow TA, Kaufman TC, Kellis M, Gelbart W, Iyer VN, et al. 2007. Evolution of genes and genomes on the Drosophil aphylogeny. Nature 450:203–218.

Danecek P, Auton A, Abecasis G, Albers CA, Banks E, DePristo MA, Handsaker RE, Lunter G, Marth GT, Sherry ST, et al. 2011. The variant call format and VCFtools. Bioinformatics 27:2156–2158.

Deng X, Gu W, Federman S, Plessis L du, Pybus OG, Faria NR, Wang C, Yu G, Bushnell B,Pan C-Y, et al. 2020. Genomic surveillance reveals multiple introductions of SARS-CoV-2 into Northern California. Science 369:582–587.

Fernandes JD, Hinrichs AS, Clawson H, Gonzalez JN, Lee BT, Nassar LR, Raney BJ, Rosenbloom KR, Nerli S, Rao AA, et al. 2020. The UCSC SARS-CoV-2 Genome Browser. Nature Genetics 52:991–998.

Grüning B, Dale R, Sjödin A, Chapman BA, Rowe J, Tomkins-Tinch CH, Valieris R, Köster J. 2018. Bioconda: sustainable and comprehensive software distribution for the life sciences. Nat Methods 15:475–476.

Hadfield J, Megill C, Bell SM, Huddleston J, Potter B, Callender C, Sagulenko P, Bedford T, Neher RA. 2018. Nextstrain: real-time tracking of pathogen evolution. Bioinformatics 34:4121–4123.

Hodcroft EB, Maio ND, Lanfear R, MacCannell DR, Minh BQ, Schmidt HA, Stamatakis A, Goldman N, Dessimoz C. 2021. Want to track pandemic variants faster? Fix the bioinformatics bottleneck. Nature 591:30–33.

Hubisz MJ, Pollard KS, Siepel A. 2011. PHAST and RPHAST: phylogenetic analysis with space/time models. Briefings in Bioinformatics 12:41–51.

Junier T, Zdobnov EM. 2010. The Newick utilities: high-throughput phylogenetic tree processing in the UNIX shell. Bioinformatics 26:1669–1670.

Lanfear R, Mansfield R. 2020. A global phylogeny of SARS-CoV-2 sequences from GISAID. Zenodo Available from: https://zenodo.org/record/3958883

Mai U, Mirarab S. 2018. TreeShrink: fast and accurate detection of outlier long branches in collections of phylogenetic trees. BMC Genomics 19:272.

Maxmen A. 2021. One million coronavirus sequences: popular genome site hits mega milestone. Nature 593:21–21.

Nicholls SM, Poplawski R, Bull MJ, Underwood A, Chapman M, Abu-Dahab K, Taylor B, Jackson B, Rey S, Amato R, et al. 2020. MAJORA: Continuous integration supporting decentralised sequencing for SARS-CoV-2 genomic surveillance.bioRxiv:2020.10.06.328328.

Paradis E, Schliep K. 2019. ape 5.0: an environment for modern phylogenetics and evolutionary analyses in R. Bioinformatics 35:526–528.

Parker J, Rambaut A, Pybus OG. 2008. Correlating viral phenotypes with phylogeny: accounting for phylogenetic uncertainty. Infect Genet Evol 8:239–246.

Rambaut A, Holmes EC, O’Toole Á, Hill V, McCrone JT, Ruis C, du Plessis L, Pybus OG. 2020. A dynamic nomenclature proposal for SARS-CoV-2 lineages to assist genomic epidemiology. Nature Microbiology 5:1403–1407.

Salemi M, Lamers SL, Yu S, de Oliveira T, Fitch WM, McGrath MS. 2005. Phylodynamic Analysis of Human Immunodeficiency Virus Type 1 in Distinct Brain Compartments Provides a Model for the Neuropathogenesis of AIDS. J Virol 79:11343–11352.

Shu Y, McCauley J. 2017. GISAID: Global initiative on sharing all influenza data – from vision to reality. Eurosurveillance 22:30494.

da Silva Filipe A, Shepherd JG, Williams T, Hughes J, Aranday-Cortes E, Asamaphan P, Ashraf S, Balcazar C, Brunker K, Campbell A, et al. 2021. Genomic epidemiology reveals multiple introductions of SARS-CoV-2 from mainland Europe into Scotland. Nature Microbiology 6:112–122.

Turakhia Y, De Maio N, Thornlow B, Gozashti L, Lanfear R, Walker CR, Hinrichs AS, Fernandes JD, Borges R, Slodkowicz G, et al. 2020. Stability of SARS-CoV-2 phylogenies.Barsh GS, editor. PLoS Genet 16:e1009175.

Turakhia Y, Thornlow B, Hinrichs AS, De Maio N, Gozashti L, Lanfear R, Haussler D, Corbett-Detig R. 2021. Ultrafast Sample placement on Existing tRees (UShER) enables real-time phylogenetics for the SARS-CoV-2 pandemic. Nature Genetics:1–8.

Wang TH, Donaldson YK, Brettle RP, Bell JE, Simmonds P. 2001. Identification of shared populations of human immunodeficiency virus type 1 infecting microglia and tissue macrophages outside the central nervous system. J Virol 75:11686–11699.

